# The Estuary Effect: Variations in Temperature and Salinity Alter *msh* Promoter Activity in *Vibrio cholerae*

**DOI:** 10.64898/2026.09.27.754871

**Authors:** Debajjyoti Basu, Gursewak Bains, Ben Ross, Andrew Michael, Joseph Alexander, Anindita Saha, Kyle A. Floyd

## Abstract

*Vibrio cholerae*, the facultative pathogen underlying cholera, naturally inhabits warm aquatic estuaries. Environmental persistence is enhanced by the ability of *V. cholerae* to colonize host reservoirs and form multicellular biofilms, causing seasonally endemic outbreaks in many tropical regions. Most toxigenic strains utilize the type IVa mannose-sensitive hemagglutinin (MSHA) pilus for host reservoir colonization and biofilm formation. Temperature and salinity can alter *V. cholerae* biofilm formation, yet their impact on MSHA production specifically remains largely unknown. Here, we utilized transcriptional reporters of predicted *msh* promoters (*msh*-P1/*msh*-P2/*msh*-P3) and functional assays, to determine temperature and salinity impacts on *msh* expression and pilus biogenesis. Under standard laboratory conditions (30°C, 1% NaCl) only *msh*-P1/P2 are active and inversely-coordinated with one another. Both *msh*-P1/P2 activity were elevated by high temperature (37°C) and low salinity (0.25%/0.5% NaCl), and reduced by low temperature (20°C/25°C) and high salinity (2%/3% NaCl). Temperature-mediated alterations in promoter activity were not immediately reflected in changes to cell-surface MSHA levels, whereas high salinity led to decreased MSHA production. Combining high temperature (37°C) and high salinity (2%/3% NaCl), attenuated the salinity-mediated reduction of *msh*-P1/P2 activity. Biofilm biomass levels were only substantially heightened at 25°C and 20°C, likely a result of no temperature-dependent changes in cell-surface MSHA, and additional temperature-controlled biofilm regulation previously described. We also found *msh*-P1/P2 promoter activity and MSHA production varies widely across toxigenic O1 and O139 serogroups despite complete sequence homology. These results shed new light on how key signals regulate MSHA pilus production to support *V. cholerae* persistence in aquatic environments.

**Importance:** Toxigenic *Vibrio cholerae* bacteria, are the causative agent of the life-threatening pandemic diarrheal disease cholera. Along with sporadic outbreaks often following catastrophic natural disasters or large-scale breakdowns of civil sanitation infrastructure, cholera is endemic to ∼50 countries across primarily Asia and Africa, driven by *V. cholerae*’s survival in brackish estuary waters. This survival is enhanced by the ability of *V. cholerae* to interact with estuarial surfaces, via the type IVa mannose-sensitive hemagglutinin (MSHA) pilus. This study identifies the impacts of key estuarial signals, temperature and salinity, on *msh* gene promoter activity and pilus production. Our results indicate that promoter activity and pilus production is reduced at lower temperatures and higher salinities, which perhaps helps to explain *V. cholerae*’s preferential habitation of warm brackish aquatic environments.

## Introduction

*Vibrio cholerae* is a natural inhabitant of aquatic environments, and a facultative human pathogen responsible for the deadly pandemic gastrointestinal disease cholera (*1*). Consumption of *V. cholerae* contaminated water or food leads to the development of severe watery diarrhea, induced via bacterial production of cholera toxin (CTX), resulting in extreme dehydration and possibly death (*1*, *2*). Large-scale cholera epidemics often follow catastrophic natural disasters or widespread civil conflicts that cause a breakdown of sanitation or water purification capabilities in regions already resource-limited (*3*). Beyond sporadic epidemics, cholera is endemic to approximately 50 countries in tropical regions of Sub-Saharan Africa and Southeast/Southwest Asia, which endure recurrent seasonal disease outbreaks estimated at approximately 2.86 million cases annually (*1*, *3*). The preferred environmental niche for *V. cholerae* is warm (∼25-30°C) brackish estuarial waters, which often coincide with high human population density, and serve as an important source of aquaculture for the population (*1*, *4*–*7*). Within this environment algae, shellfish (e.g., crustaceans and mollusks), waterfowl, protozoa, and copepods are known host reservoirs for *V. cholerae* (*8*–*19*). The ability of *V. cholerae* to survive and thrive inside such environments and reservoirs, is directly correlated with endemic seasonal cholera outbreaks within these global regions (*3*, *6*).

Persistence of *V. cholerae* within the aquatic environment is enhanced by the ability to colonize biotic host reservoirs, and form multicellular biofilm communities on both biotic and abiotic surfaces (*20*). Ingestion of environmental biofilm particles is also concomitant with exacerbated cholera disease morbidity (*7*, *21*–*25*). Biofilm formation is initiated by bacterial conversion from a free-living motile lifestyle, to a surface-associated sessile existence, mediated by attachment to a surface that induces expression of key biofilm-associated genes (*21*, *26*). There are over 200 serogroups of *V. cholerae*, encompassing both toxigenic (disease-inducing) and non-toxigenic strains, differentiated based on the structure of their lipopolysaccharide O-antigen (*27*). Toxigenic CTX-producing strains belonging to the O1 (Classical and El Tor biotypes) and O139 serogroups, are largely responsible for the current ongoing seventh cholera pandemic (*27*). For O1 El Tor and O139 serogroups, environmental surface interactions and attachment are largely driven by the type IV mannose-sensitive hemagglutinin (MSHA, **Figure 1A-B**) pilus (*26*, *28*–*34*). Type IV pili are dynamic extendable and retractile nanomachines that can facilitate surface interactions, the uptake of environmental DNA, and twitching motility (*35*). MSHA pili have been directly linked to the ability of *V. cholerae* to interact with chitinous surfaces of environmental host reservoirs, such as shellfish and zooplankton, as well as abiotic surfaces (*36*–*38*). O1 El Tor strains deficient in MSHA pilus production are highly attenuated for surface attachment and biofilm formation (*26*, *34*, *39*). Therefore, understanding the role and regulation of MSHA pili in *V. cholerae* environmental persistence, is vital to deciphering the impacts on host exposure and potential for infection.

**Figure 1.**
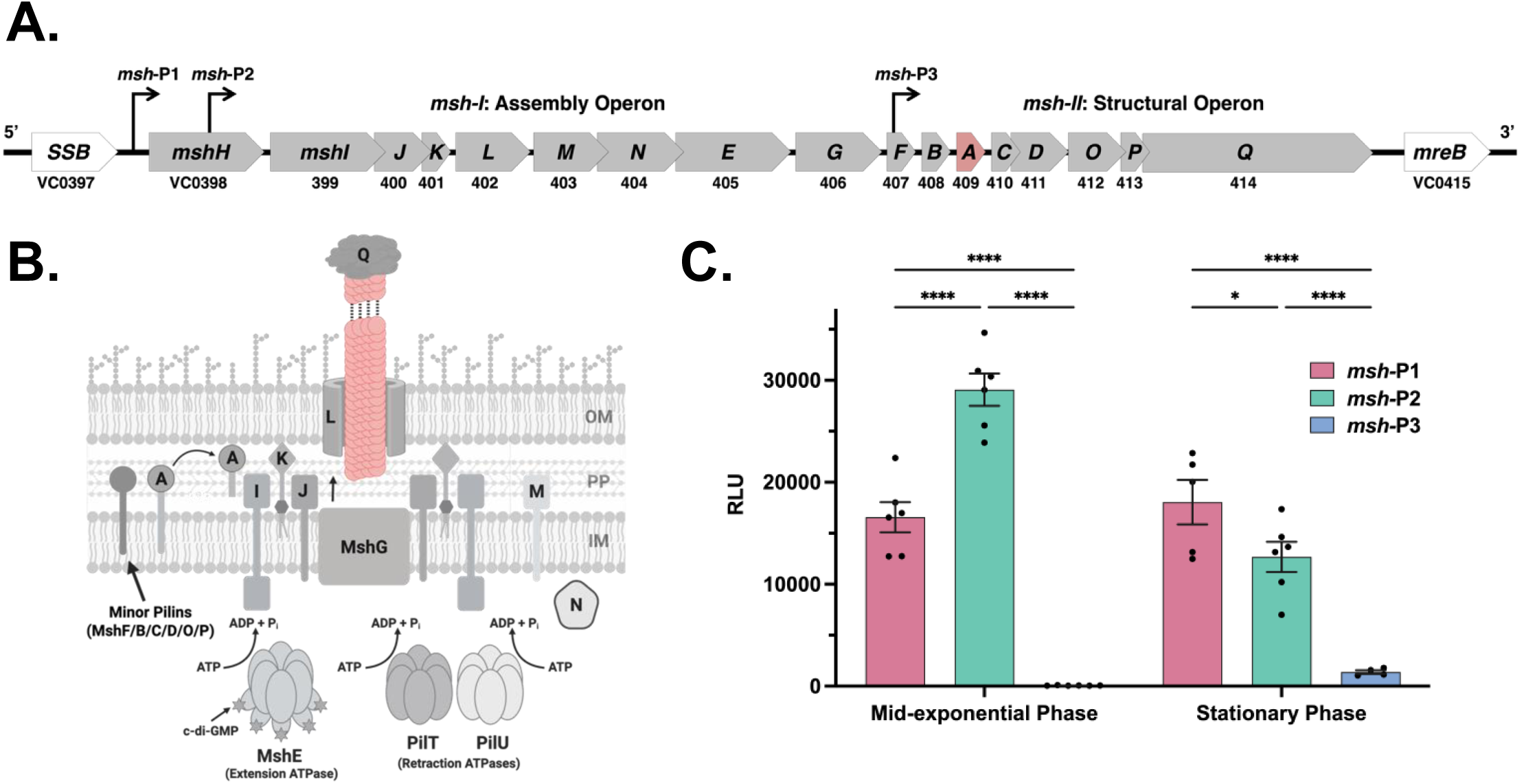
Genetic regulation of the type IVa MSHA pilus in *V. cholerae.* **(A)** Schematic representation of the predicted *msh* operons in *V. cholerae*, with reputed promoters indicated. The genes encoding the type IV MSHA pilus components are organized into two predicted operons, one containing the secretory/assembly components (*msh-I*: *mshH/I-F*), and a second containing structural components (*msh-II: mshB-Q*). Both operons are shown in schematic form with the size of the arrow indicative of the size of the coding sequence. Arrows that overlap indicate overlaps in the coding regions for respective genes. The major pilin subunit gene, *mshA*, is shown in pink. All other *msh* genes are shown in grey, with genes flanking either side of the operon(s) shown in white. This model adapted from Marsh and Taylor [1]. **(B)** Schematic of the predicted MSHA alignment/motor complex and pilus, based on homology of *msh* genes to other type IV pilus systems. **(C)** Activity of *msh*-P1 and *msh*-P2 appears inversely regulated with one another in the same growth phase. Data shown is from mid-exponential and stationary growth phases, for each of the three predicted *msh* promoters, as determined using individual pBBR*lux* transcriptional reporter assays. RLU = relative light units. Data given as mean ± SEM, from a minimum of three biological replicates. Statistical analysis: Two-way ANOVA, with Tukey correction for multiple comparisons; \**P* = 0.0342, \*\*\**P* = 0.0009, and \*\*\*\**P* < 0.0001.

MSHA pili are purportedly comprised of approximately 18 different protein subunits (**Figure 1B**); including five that constitute the dual membrane-spanning motor/assembly complex, one major pilin, five to six putative minor pilins, one adhesin, one polymerization or extension ATPase enzyme, two depolymerization or retraction ATPase enzymes, and three of unknown function (*20*). These components have been characterized as encoded by three distinct genetic operons, each serving a specific function in the assembly and operation of the pilus structure (**Figure 1A**) (*31*). The primary *msh* gene locus includes the predicted *msh-I* and *msh*-*II* operons. The *msh-I* “assembly” operon encodes genes for proteins involved in construction and function of the motor/assembly complex, and extension of pili on the cell-surface (**Figure 1A-B**); including *mshH, mshI, mshJ, mshK, mshL, mshM, mshN, mshE* (extension/polymerization ATPase)*, mshG, and mshF* (*31*). The *msh-II* “structural” operon encodes genes for proteins that collaboratively form the core structure of the pilus itself (**Figure 1A-B**); including *mshB, mshA, mshC, mshD, mshO, mshP, and mshQ* (*31*). Lastly, the third operon *pilTpilU*, located outside of the primary *msh* locus, encodes for the ATPase(s) responsible for pilus depolymerization and retraction (*40*, *41*). The *msh-I* operon has been described to have two promoter regions, one positioned upstream of *mshH* (*msh*-P1), and another internal to *mshH* (*msh*-P2); while the *msh-II* operon is described to have a single promoter internal to the final gene of the *msh-I* operon, *mshF* (*msh*-P3) (**Figure 1A**) (*31*).

Surface attachment, surface-sensing, and the corresponding response via biofilm-related gene expression are stringently controlled and regulated processes. MSHA-mediated *V. cholerae* surface attachment is regulated post-translationally by interactions of the secondary-messenger signaling molecule, 3’5’-cyclic diguanylate monophosphate (c-di-GMP) with the MshE polymerization/extension ATPase (*33*, *34*, *39*, *42*). Elevated intracellular c-di-GMP levels promote MshE-mediated extension activity, and limit pilus depolymerization/retraction to promote *V. cholerae* surface colonization (*34*). While there has been extensive study of this post-translational regulation of MSHA pilus production and dynamics, less is known about the transcriptional regulation of the system. To date, the only characterized *msh* transcriptional regulators known are VcRfaH (VC0990), VC1371, and the virulence-associated regulatory protein ToxT (VC0838) (*43*, *44*). ToxT represses *msh* gene expression through direct interactions with all three predicted *msh* promoters (*44*, *45*), while regulators VcRfaH (VC0990) and VC1371 have been observed to enhance activity of the *msh*-P1 promoter upstream of *mshH* (*43*). The regulation and response of each of these putative *msh* promoters to environmental and host signals remains largely to be elucidated.

*V. cholerae* estuarine habitats span vast geographical regions, from warm tropical regions of Southeast/Southwest Asia, Sub-Saharan Africa, the Caribbean, and Central/South America, to more temperate regions of the United States and Europe (*17*, *46*–*52*). Across this vast global circulation, differences in environmental conditions, including temperature and salinity, are likely to impact the distribution and persistence of *V. cholerae* (*53*, *54*). Within the Bay of Bengal and the Red Sea, areas that both experience endemic cholera outbreaks, surface water temperatures range between approximately 18-33°C and 20-33°C respectively, depending upon the season and the proximity to shore (*55*–*58*). By contrast, surface water temperatures tend to run lower in areas not typically experiencing endemic cholera outbreaks; such as the Gulf of Mexico which ranges between approximately 13-31°C, and the Mediterranean Sea which ranges between approximately 11-28°C (*59*, *60*). Seawater salinity varies by geographical location, water temperature, and proximity to the shoreline; but typically ranges from 3.1-3.8%, with an average of 3.5% (*4*). Likewise, the salinity of freshwater can also vary, but typically is less than 0.1% (*5*). Therefore, the salinity of estuarial environments will range somewhere between the salinity of freshwater and seawater depending upon the location within the estuary. Temperature and salinity are known to impact *V. cholerae* growth and biofilm formation, with biomass levels increasing at lower temperatures and salinity (*61*–*65*). However, the impact of temperature and salinity on *msh* gene expression and MSHA pilus production that underlies *V. cholerae* biofilm formation, represents a considerable gap in our current knowledge.

In this study, we utilize various molecular techniques to determine the impacts of temperature and salinity on the activity of the *msh*-P1/*msh*-P2/*msh*-P3 promoters, production of MSHA pili, and subsequent biofilm formation within pandemic-associated toxigenic *V. cholerae* O1 El Tor strain A1552. We found the *msh*-P1 and *msh*-P2 promoters to be highly dynamic in response to alterations in temperature and salinity, with the highest activity consistently observed from *msh*-P2. Despite previous reports (*31*, *44*), we were unable to observe activity of the predicted *msh*-P3 promoter under any condition. Whereas changes in temperature and salinity rapidly altered *msh*-P1 and *msh*-P2 promoter activity, only salinity immediately affected MSHA cell-surface pilus production, and only temperature modified 48-hour biofilm biomass levels. Comparison of A1552 with other O1 El Tor strains, O1 Classical biotypes, and O139 serogroups demonstrated complete *msh*-P1/P2 sequence homology, yet revealed intra- and inter-serogroup variation in promoter activity, MSHA pilus production, and biofilm formation.

## Results

Analysis of predicted *msh*-P1/P2/P3 promoter activity in *V. cholerae* O1 El Tor strain A1552.

To determine the functional activity of the three predicted *msh* promoters (**Figure 1A**), we generated transcriptional reporters for each promoter individually using the luminescence-based reporter plasmid pBBR*lux*. The promoter region used for *msh*-P1 is a 400 base pair (bp) fragment encompassing the region from +3 to -397 bp of the *mshH* gene (VC0398, pBBR*lux*::*msh*-P1), while the promoter region used for *msh*-P2 is a 350 bp fragment encompassing the region from -454 to -803 bp upstream of the +1 adenine of *mshI* (VC0399, pBBR*lux*::*msh*-P2), and the promoter region used for *msh*-P3 is a 350 bp fragment encompassing the region from -45 to -394 bp upstream of the +1 adenine of *mshB* (VC0408, pBBR*lux*::*msh*-P3). Promoter regions selected for *msh*-P2 and *msh*-P3 were selected based on sequences previously used by Marsh and Taylor (*31*).

Upon validation, each reporter plasmid was individually conjugated into the *V. cholerae* O1 El Tor strain A1552 for analysis of activity during mid-exponential and stationary growth phases. We previously determined that levels of the secondary messenger molecule c-di-GMP, which promotes cell-surface MSHA pilus polymerization/extension, are highest in mid-exponential phase for liquid grown shaking cultures (*66*). In mid-exponential phase, promoter activity of *msh*-P2 was 1.75-fold higher than the activity of *msh*-P1 (**Figure 1C**). This trend was inverted during stationary phase, where *msh*-P1 activity was 1.81-fold higher than *msh*-P2 (**Figure 1C**). To date, we have observed no significant activity from the predicted *msh*-P3 promoter region under any conditions. Here, there was no *msh*-P3 activity in mid-exponential phase, and inconsequential activity barely above background in stationary phase (**Figure 1C**). These data present a fascinating picture of the intricate regulation underlying *msh* gene expression and MSHA pilus production, with an inverse relationship between the activities of the *msh*-P1 and *msh*-P2 promoters, and either no or unknown condition-specific activity from *msh*-P3.

### Temperature-mediated effects on *msh* promoter activity and MSHA pilus production

Given that temperature has been shown to impact *V. cholerae* biofilm formation, and thereby environmental colonization and persistence, we next sought to determine the ability of temperature to alter *msh* gene expression and MSHA pilus production/function (*61*, *62*). To this end, we examined predicted *msh* promoter activity after shifting cultures from overnight growth at a standard temperature, to growth at lower or higher temperatures. Strains were cultured overnight under standard laboratory conditions (LB media, 1% NaCl) at 30°C (*65*), and sub-cultured the following day to mid-exponential phase at either the same (30°C), lower (25 and 20°C), or higher (37°C) temperatures. In mid-exponential phase we again observed the highest promoter activity from *msh*-P2 (**Figure 2A**). However, each promoter displayed a similar range of activity, with *msh*-P1 activity increasing 2.5-fold and *msh*-P2 activity increasing 2.8-fold respectively between 20°C and 37°C (**Figure 2A**).

**Figure 2.**
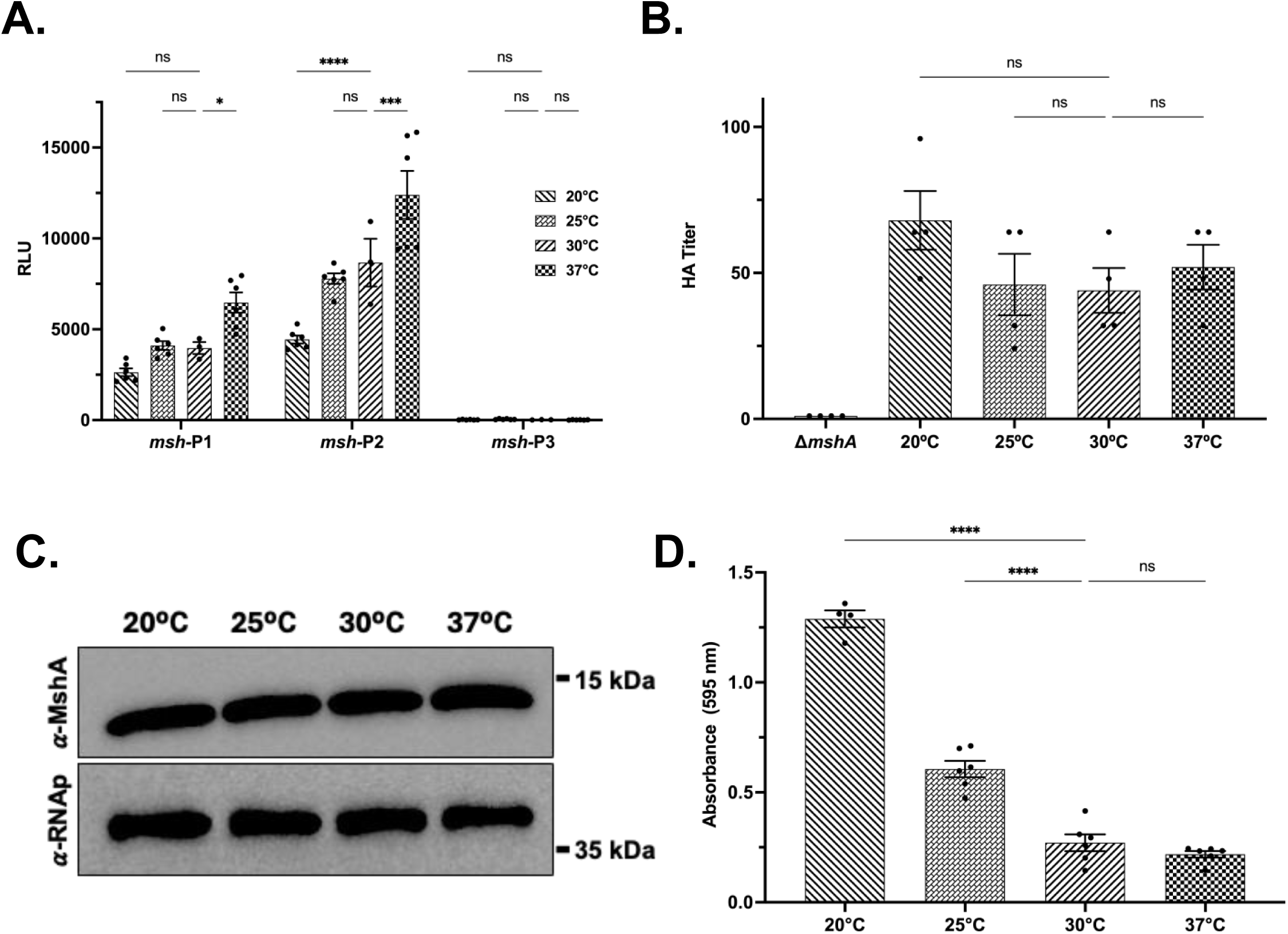
Temperature impacts on *msh* promoter activity, and MSHA pilus production. **(A)** Higher temperatures increase *msh*-P1/P2 promoter activity in mid-exponential phase (with 1% NaCl), as determined by pBBR*lux* transcriptional reporter assays. RLU = relative light units. Data given as mean ± SEM, from a minimum of three biological replicates. Statistical analysis: Two-way ANOVA with Tukey correction, comparing activity at each temperature to activity at 30°C by promoter region; \**P* < 0.0161, \*\*\**P* = 0.0003, \*\*\*\**P* < 0.0001. **(B)** Temperature does not significantly alter cell-surface MSHA levels, as determined via hemagglutination (HA) assay. The HA titer is the reciprocal of the last dilution where agglutination was observed. Data given as mean ± SEM, from a minimum of three biological replicates. Statistical analysis: (1) Indicated on graph: Ordinary One-way ANOVA comparing the mean of each temperature to one another with Dunnett’s correction for multiple comparison test; ns = not significant. (2) Not indicated on graph: Ordinary One-way ANOVA comparing the mean of each temperature to the mean of Δ*mshA* with Dunnett’s correction; [20°C: \*\*\**P* = 0.0001], [25°C: \*\**P* = 0.0047], [30°C: \*\**P* = 0.0067], [37°C: \*\**P* = 0.0017]. **(C)** Temperature does not significantly alter total cell levels of MshA major pilin subunit protein in mid-exponential phase, as determined via immunoblot. Immunoblot shown is representative of three biological replicates. **(D)** Lower temperatures increase A1552 biofilm levels at 48-hours, as determined via crystal violet assay. Data given as mean ± SEM, from a minimum of three biological replicates. Statistical analysis: Ordinary One-way ANOVA comparing the mean of each temperature to 30°C with Tukey’s correction; \*\*\*\**P* < 0.0001, ns = not significant.

For *msh*-P1, low temperature shifts from 30°C to either 25°C or 20°C resulted in no significant alteration in activity, whereas the shift to 20°C showed a 1.5-fold trend in reduction of activity; in contrast shifting to 37°C resulted in a 1.4-fold increase in activity (**Figure 2A**). For *msh*-P2, the shift to 25°C also resulted in no alteration in activity; however, shifting to 20°C resulted in a significant 2-fold decrease in activity, and shifting to 37°C resulted in a significant 1.4-fold increase in activity (**Figure 2A**). Analogous to the data reported above (**Figure 1C**), the predicted *msh*-P3 promoter region again showed no significant activity at any temperature tested (**Figure 2A**). These data suggest that temperatures below 25°C decrease *msh*-P1/P2 promoter activity, whereas temperatures above 30°C enhance activity.

To determine the rate at which these transcriptional responses are reflected at the translational and post-translational levels, we examined MSHA pilus production and function. First, we utilized a functional hemagglutination (HA) assay, which estimates cell-surface piliation based on the ability of serially-diluted bacteria to agglutinate sheep erythrocytes, where higher HA titers indicate that fewer bacteria are required for agglutination due to heightened cell-surface MSHA production in that strain or growth condition (*66*). For all HA assays, a *V. cholerae* strain lacking the gene for the MSHA major pilin subunit, *mshA* (Δ*mshA*), that does not produce cell-surface MSHA pili, was used as a negative control (**Figure 2B**). Overall, there were no significant differences in mid-exponential phase HA titers between the temperatures tested, despite each demonstrating HA titers approximately 50-fold higher than the Δ*mshA* strain (**Figure 2B**). The HA assay is limited in its ability to differentiate between more modest differences in cell-surface MSHA levels, as likely the presence of a single pilus is all that is required for erythrocyte binding. Therefore, we also examined for total cell-associated major pilin subunit, MshA, protein levels via immunoblot. Between the temperatures tested, we observed no significant difference in cell-associated MshA protein levels, further validating our HA titer observations (**Figure 2C, Figure S1A**).

As a final analysis for MSHA pilus function, and to verify that our temperature growth conditions replicated those previously observed to impact *V. cholerae* biofilm formation (*61*, *62*), we analyzed biofilm levels at each temperature via crystal violet assay (*67*). For this analysis, *V. cholerae* inoculums were grown overnight (LB media, 1% NaCl) at their respective temperature, and subsequently diluted into fresh LB media containing 1% NaCl, and seeded into PVC biofilm plates. Parallel to what was observed previously (*61*, *62*), we found biofilm levels at 48-hours were reduced as temperatures increased between 20°C to 37°C (**Figure 2D**). Biofilm levels were highest at 20°C, and were 2-fold, 5-fold, and 6-fold higher than levels at 25°C, 30°C, and 37°C respectively (**Figure 2D**). Between 25°C and 30°C biofilm levels decreased 2-fold, and there was no significant difference in biomass levels between 30°C and 37°C (**Figure 2D**). Given there were no observable differences in either HA titer or MshA protein levels, differences in 48-hour biofilm biomass are likely attributed to temperature-mediated effects on other biofilm regulatory pathways as previously observed (*61*, *62*). These data suggest that despite temperature-dependent fluctuations in *msh* promoter activity, MSHA protein abundance and function are invariant, indicating delayed or decoupled translation/post-translational processing of the changes.

### Salinity-mediated effects on *msh* promoter activity and MSHA pilus production

Brackish estuarine waters, where freshwater (salinity < 0.1%) merges with oceanwater (salinity ∼3.5%), are the preferred environmental niche of *V. cholerae* (*4*, *5*). Given that *V. cholerae* can inhabit regions anywhere along this salinity gradient, we sought to determine the impacts of varying salinity on *msh* gene expression and MSHA production/function. As with the temperature analysis, inoculums were cultured overnight under standard conditions (LB media, 1% NaCl, 30°C), and the following day sub-cultured to mid-exponential phase in fresh LB media with either 0.25%, 0.5%, 1%, 2%, or 3% NaCl (0.043, 0.086, 0.171, 0.342, 0.513 M, respectively) for evaluation of promoter activity. Both *msh*-P1 and *msh*-P2 exhibited a stepwise activity range, varying by approximately 17-fold from the highest activity at 0.25% NaCl to lowest at 3% NaCl (**Figure 3A**). Despite similar salinity-mediated alterations in activity the overall activity of *msh*-P2 was almost double that of *msh*-P1, while again the *msh*-P3 region displayed no activity (**Figure 3A**).

**Figure 3.**
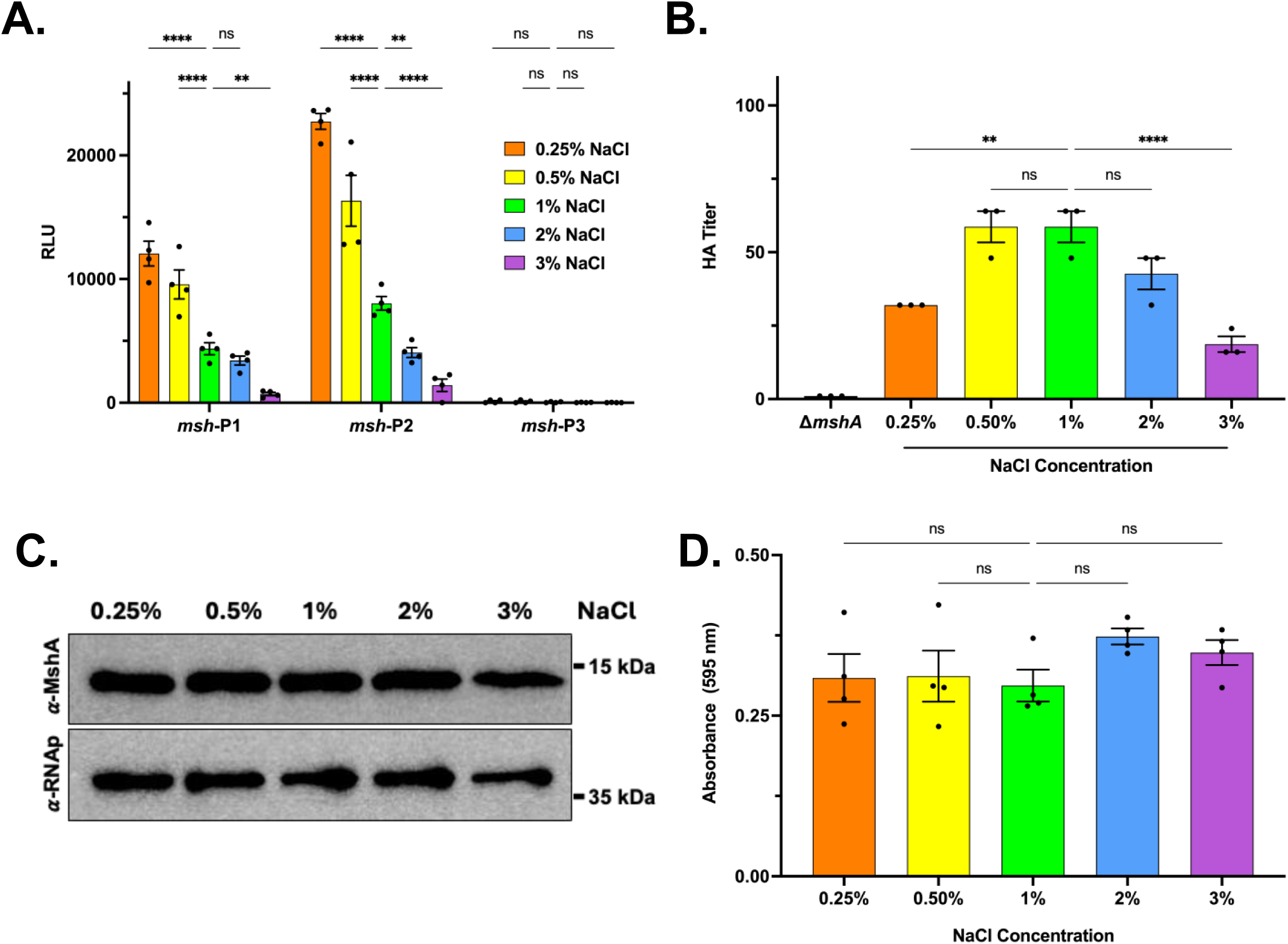
Salinity impacts on *msh*-P1/P2 promoter activity, and MSHA pilus production/function. **(A)** The promoter activity of *msh*-P1/P2 in mid-exponential phase decreases as salinity levels increase (at 30°C), as determined by pBBR*lux* transcriptional reporter assays. RLU = relative light units. Data given as mean ± SEM, from a minimum of three biological replicates. Statistical analysis: Two-way ANOVA with Tukey correction, comparing activity at each salinity to activity at 1% NaCl by promoter region; \*\**P* < 0.005, \*\*\*\**P* < 0.0001, ns = not significant. **(B)** Variations in Salinity alter cell-surface MSHA levels, as determined via hemagglutination (HA) assay. The HA titer is the reciprocal of the last dilution where agglutination was observed. Data given as mean ± SEM, from a minimum of three biological replicates. Statistical analysis: (1) Indicated on graph: Ordinary One-way ANOVA comparing the mean of each salinity to one another with Sidak’s correction; \*\**P =* 0.0022, \*\*\*\**P <* 0.00001, ns = not significant. (2) Not indicated on graph: Ordinary One-way ANOVA comparing the mean of each salinity to the mean of Δ*mshA* with Dunnett’s correction: [0.25%: \*\*\**P* = 0.0005], [0.5%: \*\*\*\**P* < 0.0001], [1%: \*\*\*\**P* < 0.0001], [2%: \*\*\*\**P* < 0.0001], [0.5%: \**P* = 0.0304]. **(C)** Salinity does not significantly alter total cell levels of MshA major pilin subunit protein in mid-exponential phase, as determined via immunoblot. Immunoblot shown is representative of three biological replicates, across multiple days. **(D)** Salinity does not significantly alter A1552 biofilm levels at 48-hours, as determined via crystal violet assay. Data given as mean ± SEM, from a minimum of three biological replicates. Statistical analysis: Ordinary One-way ANOVA comparing the mean of each salinity, with Tukey’s correction; ns = not significant.

For *msh*-P1, activity increased 2.2-fold and 2.8-fold as NaCl levels were reduced from 1% to 0.5% and 0.25% respectively (**Figure 3A**). Increasing NaCl levels from 1% to 2% did not alter *msh*-P1 activity, while increasing from 1% to 3% significantly diminished activity 6-fold (**Figure 3A**). We observed a similar pattern for *msh*-P2, where activity increased 2-fold and 2.8-fold as NaCl levels were reduced from 1%, to 0.5% and 0.25% respectively (**Figure 3A**). For *msh*-P2, activity declined 2-fold when NaCl levels were raised from 1% to 2%, and 5.6-fold when raised to 3% (**Figure 3A**). Despite *msh*-P1 and *msh*-P2 having similar trends in salinity-mediated adjustment of promoter activity, modulation of activity at higher salinities appears more significant for *msh*-P2 compared to *msh*-P1 (e.g., 1% vs 2% NaCl, **Figure 3A**), suggesting that *msh*-P2 may be more responsive to alterations in salinity than *msh*-P1.

Contrary to our observations with variable temperature, transcriptional changes resulting from salinity-induced modifications resulted in heightened translational and/or post-translational responses. HA titers were highest at 0.5% and 1% NaCl, with no difference between them (**Figure 3B**). Between 1% and 2% NaCl there was a non-significant 1.4-fold decreasing trend in HA titers, and a significant 3-fold decrease between 1% and 3% NaCl (**Figure 3B**). Interestingly, despite the highest activity of *msh*-P1/P2 occurring at 0.25% NaCl, HA titers were reduced 1.8-fold from titers at 1%, suggesting potential impacts at the translational or post-translational levels (**Figure 3B**). MshA immunoblot analysis again showed no significant differences in total cell-associated protein levels between salinities (**Figure 3C, Figure S1B**), implying possible post-translational responses at 0.25% NaCl. However, some replicates demonstrated minor highly replicate-variable decreases in MshA protein levels at 3% NaCl, and as such we are currently evolving our methods to distinguish more modest differences in protein levels between conditions. However, despite transcriptional/post-translational regulatory responses to salinity variation, there were no observable differences in biofilm levels at 48-hours (**Figure 3D**). This is similar to previous reports of no differences in 48-hour biofilms between 0.2 and 0.5 M NaCl (*64*). These data suggest *msh* gene expression is enhanced at lower salinities, and while salinity-induced transcriptional changes may begin to minorly impact MshA production at the translational level, more rapid responses appear to occur at the post-translational level to rapidly reduce cell-surface MSHA pilus presentation.

### Multifactorial effects of temperature and salinity on *msh* promoter activity and MSHA pilus production

Temperature and salinity levels can vary substantially across the estuarial niches *V. cholerae* inhabit. Above, we observed that *msh*-P1/P2 promoter activity increased as temperatures increased (**Figure 2A**), and salinity decreased (**Figure 3A**). We next sought to determine if either temperature or salinity exhibited a hierarchical effect on *msh*-P1/P2 promoter activity. Similar to our previous data, in mid-exponential phase, *msh*-P1 and *msh*-P2 activities were highest at 37°C and 0.25% NaCl, and overall activity of *msh*-P2 was higher than that of *msh*-P1 (**Figure 4, Figure S2**). For *msh*-P1, temperature effects on activity patterns were largely similar to prior observations, where *msh*-P1 activity was only significantly increased at 37°C across all salinities tested (**Figure 4A and 4C, Figure S2A**). Again, increasing salinity largely decreased *msh*-P1 activity in a somewhat stepwise manner; however, the trending and significant decreases in activity observed between 1% to 2% and 1% to 3% NaCl at 30°C were attenuated at 37°C (**Figure 4A and 4C, Figure S2A**). Trends in *msh*-P2 promoter activity were largely similar to those observed for *msh*-P1 (**Figure 4B-C, Figure S2B**). Interestingly, the significant decrease in *msh*-P2 activity observed between 1% and 2% and 1% and 3% NaCl at 30°C, were also attenuated at 37°C as was observed for *msh*-P1 (**Figure 4B and 4C, Figure S2B**). This suggests that high temperature-mediated enhancement (>30°C) of *msh*-P1/P2 activity supersedes the suppression induced by increased salinity (2%, 3% NaCl).

**Figure 4.**
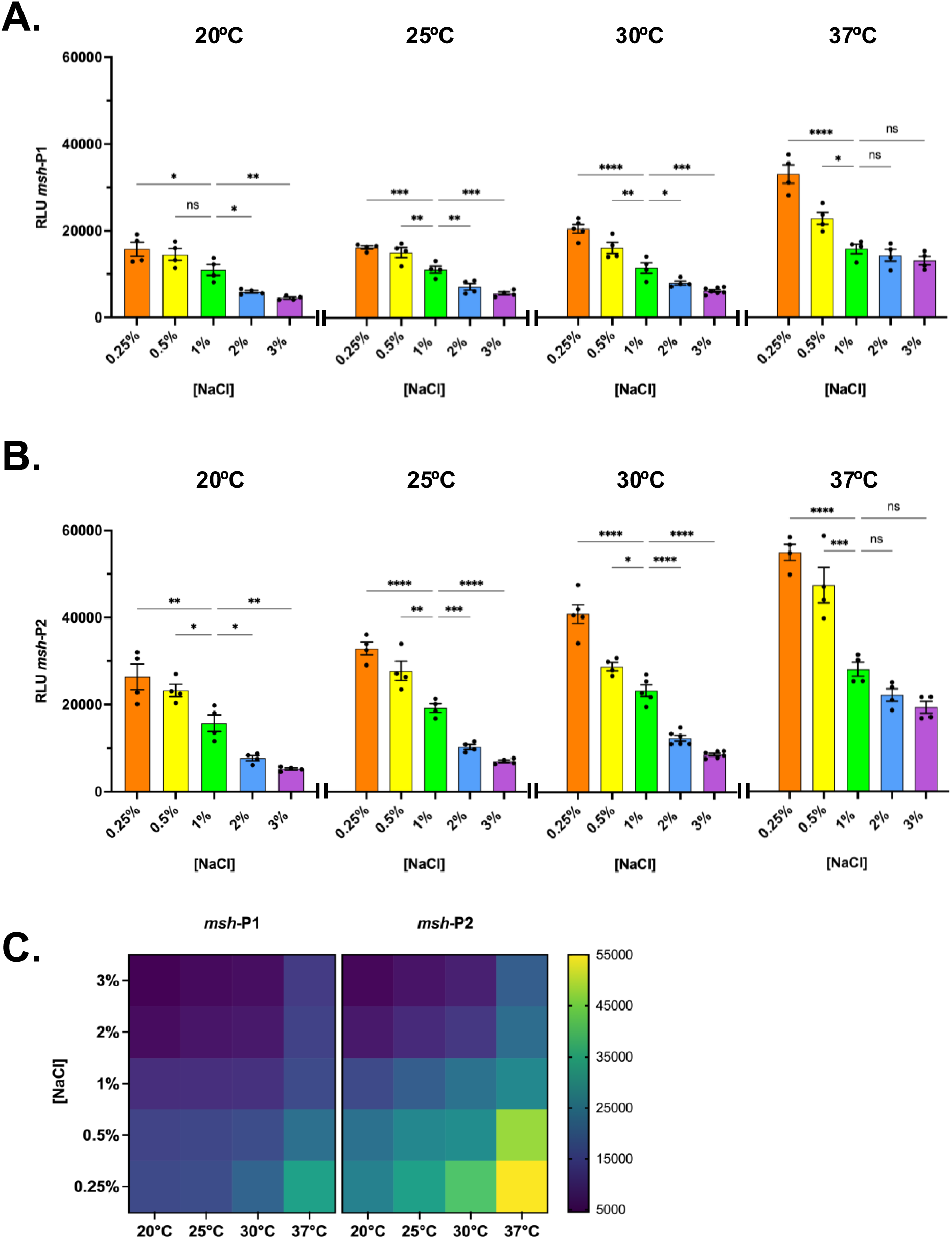
Compounded impacts of temperature and salinity on *msh*-P1/P2 promoter activity. **(A-B)** RLU = relative light units. Data given as mean ± SEM, from a minimum of three biological replicates. **(A)** Promoter activity of *msh*-P1 in mid-exponential phase. Statistical analysis: Ordinary one-way ANOVA with Dunnett’s correction, comparing activity at each salinity to 1% NaCl within temperature; [20°C: \**P* < 0.05, \*\**P* < 0.005], [25°C: \*\**P =* 0.008, \*\*\**P* < 0.001], [30°C: \**P =* 0.0461, \*\**P* = 0.006, \*\*\**P* = 0.0006, \*\*\*\**P* < 0.0001], [37°C: \**P =* 0.0124, \*\*\*\**P* < 0.0001], ns = not significant. **(B)** Promoter activity of *msh*-P2 in mid-exponential phase. Statistical analysis: Ordinary one-way ANOVA with Dunnett’s correction, comparing activity at each salinity to 1% NaCl within temperature; [20°C: \**P* < 0.03, \*\**P* < 0.002], [25°C: \*\**P* = 0.0011, \*\*\**P* < 0.001], [30°C: \**P =* 0.0195, \*\*\*\**P* < 0.0001], [37°C: \*\*\**P =* 0.0001, \*\*\*\**P* < 0.0001], ns = not significant. **(C)** Data from (A-B) re-plotted as a heatmap, based on RLU intensity values.

### Serogroup-, biotype-, and strain-specific variation in *msh* promoter activity and MSHA pilus production

Perhaps our most interesting observation from this work was the lack of activity from the putative *msh*-P3 promoter region that had been previously described (*31*, *44*). Original analysis was conducted in the O1 El Tor Inaba strain C6706 from Peru, which was obtained from the U.S. Centers for Disease Control (*31*, *68*); while our analysis was performed with the O1 El Tor Inaba strain A1552 (originally 92A1552) obtained from California state health authorities, and linked to a cholera outbreak in the 1990s also in Peru (*69*, *70*). Therefore, we decided to compare the activity of the predicted *msh* promoter regions and resulting MSHA pilus production/function between the commonly used O1 El Tor strains A1552, C6706, and E7946 (*71*); as well as other O1 biotypes (O1 Classical strains O395(1) and O395(2) (*72*)), and other serogroups (O139 strains MO10 and MO45 (*73*)). Only the O1 El Tor and O139 serogroups have previously been identified to produce MSHA pili (*29*). DNA sequences of the *msh*-P1/P2/P3 promoter regions used to generate our transcriptional reporters, share 100% sequence homology between O1 El Tor and O1 Classical biotypes, and with the O139 serogroup (**Figure S3**). In fact, O1 El Tor and O139 serogroups share 100% sequence homology within the *msh* locus, while O395 shares 99.977% homology (three single nucleotide variants, one nucleotide deletion) with both the O1 El Tor biotype and the O139 serogroup (**Figure S3**).

For O1 El Tor strain A1552 in mid-exponential phase at 30°C (LB media, 1% NaCl), similar to our previous results (**Figure 1C**); *msh*-P2 activity was 1.68-fold higher than that of *msh*-P1, and there was no activity of *msh*-P3 (**Figure 5A**). Overall, there was no observable *msh*-P3 activity within any of the strains, biotypes, or serogroups tested (**Figure 5A**). The dynamics of *msh*-P1/P2 activity were similar across O1 El Tor strains; with *msh*-P2 activity 1.6-fold higher than *msh*-P1 in E7946, and 1.8-fold higher in C6706 (**Figure 5A**). However, overall *msh*-P1/P2 activity was significantly lower in both E7946 and C6706 (∼2.4-fold), compared to A1552 (**Figure 5A**). These differences in *msh*-P1/P2 promoter activity resulted in a parallel decrease in MSHA pilus production. HA titers were reduced 2.7-fold for E7496, and 1.9-fold for C6706, compared to A1552 (**Figure 5B**), despite no difference in total cell-associated MshA protein levels between strains (**Figure 5C, Figure S1C**). This suggests that the differences in HA titer may result from differences in post-translational rather than transcriptional regulation. Despite higher levels of *msh*-P1/P2 activity and pilus production in A1552 compared to E7946 and C6706, biofilm biomass levels at 48-hours in A1552 were no different from those of E7946, and diminished 3.6-fold compared to C6706 (**Figure 5D**). O1 El Tor strain C6706 displayed the highest biofilm biomass levels of all the *V. cholerae* biotypes, serogroups, and strains tested, with levels approximately 3.4-fold higher than other O1 El Tor strains, approximately 11-fold higher than O1 Classical biotypes, and 1.6-fold / 4.7-fold higher than O139 serogroup strains MO10 and MO45 respectively (**Figure 5D**).

**Figure 5.**
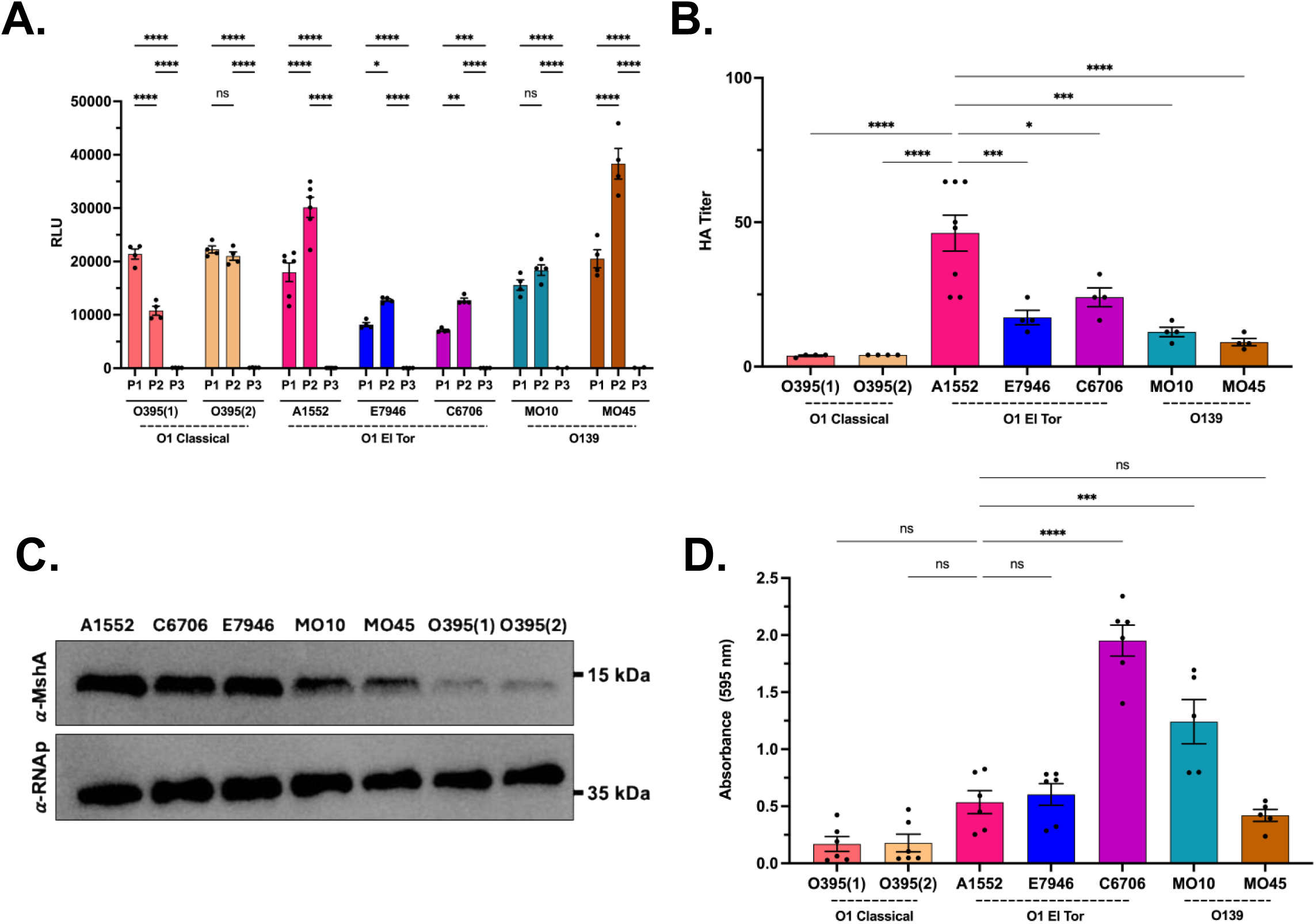
*msh* promoter activity and MSHA pilus production/function differs across O1 Classical, O1 El Tor, and O139 *V. cholerae* strains. We analyzed *V. cholerae* O1 Classical strains O395(1) and O395(2); O1 El Tor strains A1552, E7946, and C6706; O139 strains MO10 and MO45. (A) The promoter activity of *msh*-P1/P2 varies across *V. cholerae* strains, biotypes, and serogroups (at 30°C, 1% NaCl) as determined by pBBR*lux* transcriptional reporter assays. RLU = relative light units. Data given as mean ± SEM, from a minimum of three biological replicates. (1) Indicated on graph: Statistical analysis: Comparisons made between promoter activity within each strain, via Two-way ANOVA with Tukey’s correction; \**P* = 0.0263, \*\**P* = 0.0052, \*\*\*\**P* < 0.0001, ns = not significant. (2) Not indicated on graph – *msh*-P1: Statistical analysis: Comparisons made between promoter activity of each strain and O1 El Tor strain A1552, via Two-way ANOVA with Dunnett’s correction; [O395(1): not significant], [O395(2): \**P* = 0.0484], [E7946: \*\*\*\**P* < 0.0001], [C6706: \*\*\*\**P* < 0.0001], [MO10: not significant], [MO45: not significant]. (3) Not indicated on graph – *msh*-P2: Statistical analysis: Comparisons made between promoter activity of each strain and O1 El Tor strain A1552, via Two-way ANOVA with Dunnett’s correction; All strains vs A1552, \*\*\*\**P* < 0.0001. (4) Not indicated on graph – *msh*-P3: Statistical analysis: Comparisons made between promoter activity of each strain and O1 El Tor strain A1552, via Two-way ANOVA with Dunnett’s correction; All strains vs A1552, not significant. **(B)** Cell-surface MSHA levels by *V. cholerae* strain, as determined via hemagglutination (HA) assay at 30°C, 1% NaCl. The HA titer is the reciprocal of the last dilution where agglutination was observed. Data given as mean ± SEM, from a minimum of three biological replicates. Statistical analysis: Comparisons made between promoter activity of each strain and O1 El Tor strain A1552, via ordinary One-way ANOVA with Tukey’s correction; \**P =* 0.0161, \*\*\**P* < 0.001, \*\*\*\**P* < 0.0001. Comparisons that are “not significant” are not shown, to reduce the complexity of the graph. **(C)** Total cellular MshA protein levels by strain, as determined via immunoblot. Immunoblot shown is representative of two biological replicates, across multiple days. **(D)** Levels of biofilm formation at 48-hours (25°C, 1% NaCl), as determined via crystal violet assay, differ among *V. cholerae* strains. Data given as mean ± SEM, from a minimum of three biological replicates. Statistical analysis: Comparisons made between biofilm biomass levels of each strain and O1 El Tor strain A1552, via ordinary One-way ANOVA with Dunnett’s correction; \*\*\**P =* 0.005, \*\*\*\**P* < 0.0001, ns = not significant.

Between the O1 El Tor and O1 Classical biotypes, both O395(1) and O395(2) strains demonstrated reduced *msh*-P2 activity (∼1.4-fold) compared to A1552 in mid-exponential phase, while *msh*-P1 activity was only slightly but not significantly enhanced (**Figure 5A**). Intriguingly, O1 Classical *in situ msh*-P1/P2 promoter dynamics differed from those observed in the O1 El Tor biotype and O139 serogroup. Strain O395(1) exhibited transposed *msh*-P1/P2 dynamics compared to A1552, where *msh*-P1 activity was 2-fold higher than that of *msh*-P2, while strain O395(2) exhibited no difference between *msh*-P1 and *msh*-P2 activity as *msh*-P2 activity was 1.9-fold higher than O395(1) (**Figure 5A**). Both O1 Classical strains exhibited negligible HA titers and cell-associated MshA protein levels, and a subsequent reduction in biofilm levels compared to O1 El Tor biotype and O139 serogroup strains; with no observable differences in these phenotypes between the two O395 strains (**Figure 5B-D, Figure S1C**).

Within the O139 serogroup, strain MO45 displayed *msh*-P1/P2 promoter dynamics inversely proportional to those of O1 El Tor strains, and demonstrated the highest overall *msh*-P2 activity (1.3-fold higher than A1552 *msh*-P2 activity, **Figure 5A**). Promoter dynamics in O139 strain MO10 exhibited a slight trend towards similarity with those in O1 El Tor, however statistically there were no differences between *msh*-P1 and *msh*-P2 activity (**Figure 5A**). Despite MO45 having the highest level of *msh*-P2 activity in mid-exponential phase, HA titers for both MO45 and MO10 were lower than most O1 El Tor strains, and marginally above the negligible levels in the O1 Classical strains (**Figure 5B**). This suggests a substantial reduction in cell-surface MSHA pilus production in the O139 serogroup compared to O1 El Tor. This is supported by the observation of reduced total cell-associated MshA protein levels, in both MO10 and MO45 compared to the O1 El Tor serogroup (**Figure 5C, Figure S1C**). This reduction in MSHA pilus production likely results from both transcriptional and translational/post-translational changes in MO10, given the reduction in *msh*-P2 activity, HA titers, and MshA protein. However, MO45 is a more interesting case, as *msh*-P2 activity is elevated compared to A1552, yet there is still a substantial reduction in HA titers and MshA protein levels. This suggests that, in MO45, regulation of MSHA pilus production likely occurs at the translational and/or post-translational level rather than the transcriptional level. Notwithstanding the decrease in MSHA pilus production, both O139 strains exhibited robust biofilm forming abilities (**Figure 5D**). MO10 had the second highest biofilm biomass levels at 48 hours amongst all strains, which were 2.3-fold higher than O1 El Tor strain A1552 (**Figure 5D**). MO45 demonstrated a 3-fold less biofilm than MO10 at 48-hours, which was equivalent to levels in O1 El Tor strains A1552 and E7946 (**Figure 5D**). These differences in MSHA production dynamics, highlight considerable diversity in MSHA pilus regulation across, and even within, *V. cholerae* biotypes, serogroups, and strains.

## Discussion

Toxigenic O1 (El Tor and Classical) and O139 *V. cholerae* serogroups, are frequently found living in freshwater and brackish water environments (*7*). *V. cholerae* survival and persistence within these estuarial environments is enhanced by their ability to interact with biotic reservoir hosts, and to form sessile multicellular biofilm communities. While the influence of MSHA pili on environmental surface interactions is well-known (*26*, *28*–*34*), the role of environmental signals in regulating *msh* gene expression and pilus production is less understood. Here, we sought to determine the impacts of the key environmental signals temperature and salinity on *msh* promoter activity and MSHA pilus production in *V. cholerae* O1 El Tor strain A1552. Using luminescence-based transcriptional reporters for the three predicted promoter regions of the *msh* locus (*msh*-P1, *msh*-P2, *msh*-P3), we determined that *msh* gene expression is primarily coordinated by the *msh*-P1 and *msh*-P2 promoters. The activity of these two promoters is inversely connected, with the highest activity coming from *msh*-P2 during mid-exponential growth when MSHA pilus production is highest. Conversely, in stationary growth phase where MSHA pilus production is reduced, *msh*-P1 activity was higher than *msh*-P2. This suggests that *msh*-P2 is the key promoter driving gene expression at the *msh* locus. This is consistent with *mshH* having no previously described role in MSHA pilus production or function, as it encodes an *Escherichia coli csrD* homolog that control production of small regulatory RNAs of the carbon storage regulator gene A (*csrA*) (*74*). Analysis of these dynamics across other *V. cholerae* O1 El Tor strains, the O1 Classical biotype, and the O139 serogroup, demonstrated that *msh*-P1/*msh*-P2 inverse promoter dynamics were conserved across MSHA-producing O1 El Tor and O139 serogroups, yet were transposed or altered in the MSHA-deficient O1 Classical biotype.

Despite previous reports (*31*, *44*), we failed to detect any promoter activity from the *msh*-P3 region across all strains, biotypes, and serogroups. Given that the *msh*-P3 region has been shown to contain functional consensus binding sequences for the virulence-associated transcriptional regulator ToxT, it is possible that the *msh*-P3 region represents a repressor site, rather than an active promoter requisite for expression of a separate *msh*-II operon (*44*). ToxT also has functional consensus binding sequences within both the *msh*-P1 and *msh*-P2 promoter regions (*44*). This possible negative regulatory function of *msh*-P3 could serve as a fail-safe mechanism to prevent structural pilin gene transcription (including the *mshA* major pilin) during host infection, in the event that transcription from the active *msh*-P1/P2 promoters has failed to cease. Based on our observations to date, we hypothesize that there is likely a single *msh* operon (*msh[H]IJKLMNEGFBACDOPQ*) regulated primarily by *msh*-P2, and secondarily by *msh*-P1; resulting in two possible mRNA transcripts (*msh1*: *mshH-Q*, *msh2*: *mshI-Q*), as opposed to two independently regulated operons. Ongoing studies in the lab are refining the *msh* genetic organization, including analysis of the *msh* transcriptional start sites and resulting mRNA transcripts, to address this hypothesis.

Estuaries are also home to many species of algae, shellfish (e.g., crustaceans and mollusks), waterfowl, protozoa, and copepods that are known host reservoirs for *V. cholerae* (*8*–*19*). The ability of *V. cholerae* to survive and thrive in such environments and reservoir hosts is directly correlated with endemic cholera disease, with shellfish being among the primary contaminated food sources resulting in human exposure to *V. cholerae* (*3*, *6*). MSHA pili are directly linked to interactions with abiotic surfaces, and chitin-containing biotic reservoir hosts in estuarine environments (*36*–*38*). MSHA pili are also highly conserved across other pathogenic *Vibrio* species, and are also tied to their ability to persist within the same estuarine environments and infect the human host (*75*). This includes the necrotizing fasciitis-causing species *V. vulnificus* and *V. parahaemolyticus*, which demonstrate a significantly higher mortality rate than that of *V. cholerae* (*75*). Therefore, understanding the signals and mechanisms that regulate *msh* gene expression and MSHA pilus biogenesis, is of vital global public health importance.

In regions of Sub-Saharan Africa and Southeast/Southwest Asia afflicted with endemic seasonal cholera epidemics (*3*), surface oceanwater temperatures can range from as low as 18°C during the coldest parts of the year, to as high as 33°C during the warmest periods when cholera cases peak (*55*–*58*). The salinity of freshwater is typically < 0.1%, while the salinity of oceanwater can be as high as approximately 3.5%; thus, estuarial salinity will vary along this 0.1-3.5% gradient based on proximity to the freshwater or oceanwater source (*4*, *5*). Here, we found that individually lower temperature (20°C, 25°C) and higher salinity (2%, 3% NaCl) significantly reduced *msh*-P1 and *msh*-P2 promoter activity. Simultaneous variation of salinity at different temperatures, largely demonstrated *msh*-P1/P2 activity trends similar to the individual trends. However, the reduction in promoter activity observed at 2% and 3% NaCl compared to 1% NaCl at 30°C, was negated when the temperature was increased to 37°C. This implies that high temperature may exert a dominant regulatory effect over those induced by high salinity. This dominance could play an important role during *V. cholerae* interactions with the host, as temperatures in the intestinal lumen average approximately 36.96°C, slightly warmer than that of blood at approximately 36.69°C (*76*). Rising global ocean temperatures resulting from the impacts of climate change, also threaten to expand the colonization range of *V. cholerae* and other pathogenic *Vibrio* species (*77*).

Despite alterations in *msh*-P1/P2 promoter activity with both signals, cellular levels of the MshA major pilin subunit were generally unchanged, and only heightened salinity (2%, 3% NaCl) resulted in an immediate decrease in cell-surface MSHA pilus levels at 30°C. This suggests a delay in imparting temperature- and salinity-mediated alterations in *msh* expression to changes at the translation and post-translational levels, with the exception that high salinity (2%, 3% NaCl) triggers a rapid post-translational response to minimize cell-surface pilus presentation. We speculate that this rapid post-translational diminishment of cell-surface MSHA pili at higher salinities could, in part, help to explain the preferential habitation of *V. cholerae* within estuarial environments. Higher MSHA production at lower salinity levels could also further impact human infection, given that NaCl levels in the lumen of the small intestines are typically isotonic to those in blood plasma at approximately 0.14 M (∼0.8%), in the range in which we find elevated promoter activity and cell-surface presentation (*78*).

The ongoing seventh cholera pandemic, which began in ∼1961, has been dominated by toxigenic O1 El Tor and O139 *V. cholerae* serotypes (*7*). Compared to the O1 Classical serotype associated with the sixth pandemic (1899-1923), O1 El Tor and O139 *V. cholerae* serotypes produce MSHA pili (*7*). We found *msh*-P1/P2 dynamics to be similar across O1 El Tor and O139 strains, yet different from those in O1 Classical strains. These altered dynamics could result from evolutionary changes in response to the lack of MSHA pilus production, or given that *msh*-P2 appears to be the most crucial *msh* promoter, could be a contributing factor resulting in their lack of MSHA production. Whichever scenario is true, we speculate that these differential dynamics in O1 Classical strains has contributed to the dominance of O1 El Tor and O139 serogroups in the current pandemic, over O1 Classical and non-O1 serogroups. Despite similar patterns in promoter dynamics, we found that overall promoter activity, cell-surface MSHA pilus levels, MshA pilin protein production, and biofilm formation vary widely not only between but also within O1 El Tor and O139 serotypes. Most intriguingly, these variations occur despite 100% sequence homology across the entirety of the *msh* locus between O1 El Tor and O139 strains; with the only differences across the *msh* locus being three single nucleotide polymorphisms and one nucleotide deletion in O1 Classical, compared to O1 El Tor and O139 strains. These differences in *msh*-P1/P2 activity, MSHA production, and biofilm formation demonstrates a high level of diversity in MSHA pilus regulation across *V. cholerae* bio- and serotypes. Ongoing work in the lab, seeks to leverage this diversity for the identification of additional *msh* transcriptional, translational, and/or post-translational regulatory pathways.

## Materials and Methods

### Bacterial Strains, Media, and Growth Conditions

Bacterial strains used in this study are listed in **Table S1**. Unless specified otherwise, all analyses were performed in the *V. cholerae* O1 El Tor strain A1552. *V. cholerae* and *Escherichia coli* strains were grown at 30 and 37°C, respectively, with 200 rpm shaking in Luria-Bertani (LB) media. Briefly, overnight cultures were prepared by inoculating five single colonies from LB-agar plates into 5 mL LB media. Overnight cultures were next diluted 1:200 into fresh 5 mL LB media, and grown to mid-exponential phase (OD_600_ ∼0.3-0.4) for analysis unless stated otherwise. Strains containing pBBR*lux* reporter plasmids were grown in the presence of 5 µg/mL (*V. cholerae*), 20 µg/mL (*E. coli*) chloramphenicol (Cm), or 100 µg/mL ampicillin (Amp) with 5 µg/mL Cm. For testing the impacts of salinity, differential LB media were generated containing 20 g/L tryptone, 10 g/L yeast extract, supplemented with 2.5, 5, 10 (Normal LB concentration), 20, 30 g/L NaCl, and the pH was adjusted to 7.5 with 10 N NaOH or 6 N HCl. For testing the impacts of temperature, cultures were grown at either 20, 25, 30, or 37°C with 200 rpm shaking.

### Generation of Transcriptional Reporter Plasmids

All plasmids used in this study are listed in **Table S1**. Luminescence-based transcriptional reporter plasmids were constructed using the pBBR*lux* plasmid containing the *luxABCDE* operon, digested with the restriction endonucleases SacI and BamHI (New England Biolabs). Primers (**Table S2**) were designed to amplify each of the predicted promoter regions of the MSHA operon(s) - specifically, *msh-*P1, *msh-*P2, and *msh-*P3, based on the promoter boundaries outlined previously (*31*, *44*). Each amplified promoter region was ligated into the pBBR*lux* plasmid upstream of *luxABCDE* using Hifi DNA Assembly Master Mix (New England Biolabs), and then transformed into chemically competent *E. coli* strain S17-1λpir for conjugation into *V. cholerae*; generating plasmids pBBR*lux*::*msh*-P1, pBBR*lux*::*msh*-P2, and pBBR*lux*::*msh*-P3. Due to intrinsic CM resistance of O139 strains MO10 and MO45, an ampicillin-resistance cassette was amplified with its constitutive promoter from plasmid pMMB67EH, and cloned into pBBR*lux*::*msh*-P1/P2/P3 plasmids between KpnI and PacI restriction sites near the Cm cassette; generating plasmids pBBR*lux_amp*::*msh*-P1, pBBR*lux_amp*::*msh*-P2, and pBBR*lux_amp*::*msh*-P3. All plasmids were verified to contain the correct promoter sequence via whole plasmid sequencing (Plasmidsaurus). Once verified, reporter plasmids were conjugated into the respective *V. cholerae* strain, and selection was made using LB-agar plates containing 5 µg/mL Cm + 100 µg/mL rifampicin, or 100 µg/mL Amp + 5 µg/mL Cm + 100 µg/mL rifampicin (Rif). Presence of the appropriate plasmid was confirmed via colony PCR before 50% glycerol freezer stocks were prepared.

### Promoter Activity Assays

Strains containing individual reporter plasmids were streaked onto LB-agar plates containing 5 µg/mL Cm and incubated statically at 30°C overnight. Overnight cultures were prepared by picking five single colonies and inoculating them into 5 mL of LB broth containing 5 µg/mL of Cm, which were then incubated at 30°C with 200 rpm shaking. A 1:200 dilution of the overnight culture was then diluted into 5 mL of their respective growth media containing 5 µg/mL Cm and incubated at their indicated temperature with 200 rpm shaking, until they reached mid-exponential phase (OD_600_ ∼0.3 - 0.4) as determined by a spectrophotometer (Eppendorf), unless otherwise stated. Levels of luminescence were then determined using an Infinite 200 Pro plate reader (Tecan). Assays were repeated with a minimum of three biological replicates. Data was plotted and statistical analysis performed using GraphPad Prism 11.

### Hemagglutination Assay

All strains analysed via hemagglutination (HA) were grown to mid-exponential phase (in their respective media and temperature) as described above, without chloramphenicol. HA assays were then performed as previously described (*66*). Defibrinated sheep blood was used for the assays (Hardy Diagnostics). Assays were repeated with a minimum of three biological replicates. Data was plotted and statistical analysis performed using GraphPad Prism 11.

### Immunoblot for Total Cellular MshA Protein Levels

For analysis of total cellular MshA major pilin subunit protein levels, cultures were grown to mid-exponential phase as described above with their respective media/temperature conditions. Immunoblotting was performed as previously described (*66*). Briefly, once cultures reached mid-exponential phase, 2 mL was centrifuged at 6000 x g for 5 minutes. The supernatant was discarded; pellets were resuspended in 300 µL of 2% sodium dodecyl sulfate (SDS) and boiled at 100 °C for 10 minutes. Total protein levels of each sample was then determined via BCA assay (Thermo Scientific). For SDS-PAGE, for each sample, 10 µg of protein in a final volume of 25 µL was loaded onto a 16% 1.5mm SDS-PAGE gel, and run at 90 V for 10 minutes and then 120 V until the dye front was just off of the gel. Proteins were then transferred to a 0.2 µm PVDF blotting membrane (Cytiva) using a Trans-Blot Turbo transfer system (Bio-Rad) at 25 V – 1.3 A for 45 minutes. Membranes were then blocked overnight in 1x PBST containing 5% non-fat milk. After blocking, membranes were incubated with primary rabbit polyclonal anti-MshA antibody (generated by GenScript) for one hour, washed twice with 1x PBST for 5 minutes each, incubated with a secondary goat anti-rabbit HRP-conjugated secondary antibody (Invitrogen) for another hour, again washed twice with 1x PBST for 5 minutes each, and visualized using SuperSignal Pico chemiluminescent substrate (Thermo Scientific) with a ChemiDoc MP system (Bio-Rad). Following image acquisition, membranes were stripped with stripping buffer (15 g/L glycine, 1 g/L SDS, 10 mL /L tween 20, pH 2.2 with 6 N HCl), re-blocked overnight, and analysis repeated using a murine polyclonal anti-*E. coli* RNA polymerase alpha antibody (BioLegend) and goat anti-mouse HRP-conjugated secondary antibody (Invitrogen) for a loading control. Immunoblots were repeated with a two to three biological replicates. Images were visualized using the Image Lab Touch Software (Bio-Rad).

### Crystal Violet Biofilm Assay

All strains analysed via crystal violet biofilm assay (*67*) were grown overnight with 200rpm shaking in their respective media and/or temperature conditions as described above, without chloramphenicol. After overnight culture, strains were diluted 1:200 in their respective media, 200 µL of each culture was plated into a 96-well plate PVC plate (Corning) and incubated statically for 48 hours at their respective temperature. After 48 hours, the cultures were removed from the 96-well plate, plates were then washed with deionized water to remove non-adherent bacteria and stained with 250 µL of 1% crystal violet for 15 minutes. Afterwards, the crystal violet was removed, plates were washed three times with deionized water, and allowed to air dry. Once the plates were completely dry, 250 µL of 96% ethanol was added to the wells to solubilize the crystal violet, and then 200 µL of each sample was transferred into a flat-bottom 96-well plate. The OD at 595 nm was then determined using an Infinite 200 Pro plate reader (Tecan). Assays were repeated with a minimum of three biological replicates. Data was plotted and statistical analysis performed using GraphPad Prism 11.

### Sequence Alignment and Analysis of the *msh* Locus Across Strains

Contiguous genomic sequences spanning the *msh* locus (*mshH*–*mshQ*, plus flanking *csgD*, *ssb*, and *mreB*) were obtained from complete chromosome assemblies of five *Vibrio cholerae* biotypes, serogroups, and strains (O1 El Tor A1552, GenBank CP028894.1; O1 El Tor E7946, CP024162.1; O1 El Tor C6706, CP157384.1; O1 Classical O395, CP045719.1; O139, ENA LT992486.1) by extracting the homologous region identified via BLASTn (v2.12.0+, NCBI). Sequences were aligned using MAFFT v7.505 (FFT-NS-2, default settings). Pairwise nucleotide identity was calculated across un-gapped alignment columns for each sequence pair. Sliding-window identity (200 bp window, 20 bp step) and per-column variant calls (substitutions and indels relative to the majority-consensus base) were computed in Python (Biopython, NumPy). Figures were generated with Matplotlib. *AI disclosure*: Sequence alignment, identity/variant calculations, and **Figure S3** generation were performed with the assistance of Claude (Anthropic, Claude Sonnet 5).

## Acknowledgments

The authors wish to thank Drs. Fitnat Yildiz (University of California – Santa Cruz) and Ankur Dalia (Indiana University) for supplying *Vibrio cholerae* strains. This research was enabled by funding generously provided by: Illinois State (ISU) University College of Arts and Sciences, School of Biological Sciences Start-up Funds to Dr. Kyle Floyd; ISU Office of Student Research FIREbird Grant to Ben Ross; ISU Phi Sigma Honor Society Robert D. Weigel Grants to Ben Ross and Debajjyoti Basu; and the National Institutes of Health, National Institute of Allergy and Infectious Diseases Grant 1R15AI185921-01 to Dr. Kyle Floyd.

## Figure Legends

**Figure S1.**
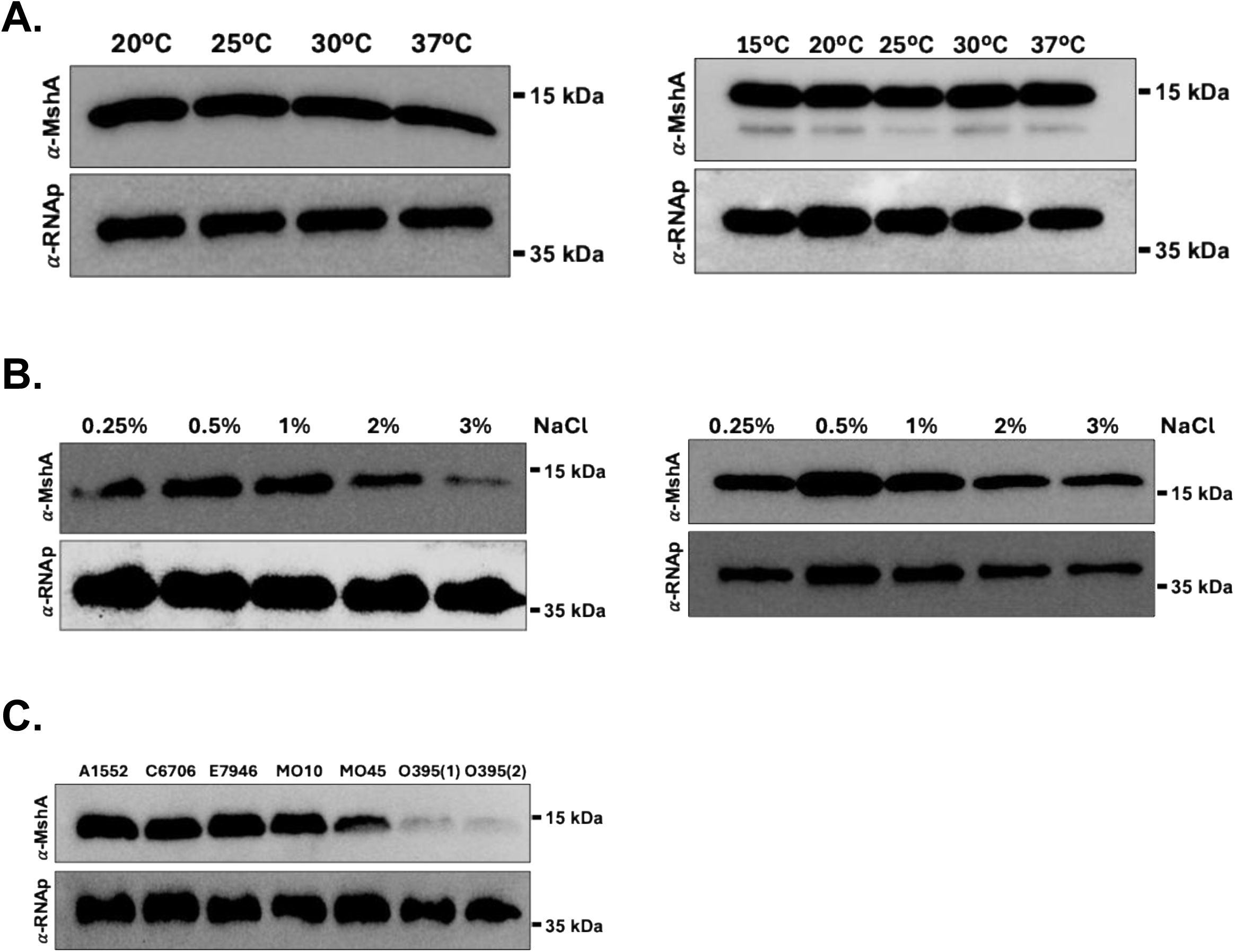
Additional biological replicate immunoblots for total cell-associated MshA protein levels, by condition. **(A**) Biological replicates two and three for immunoblot of MshA protein levels by temperature. (**B)** Biological replicates two and three for immunoblot of MshA protein levels by salinity. **(C)** Biological replicate two for immunoblot of MshA protein levels across strains.

**Figure S2.**
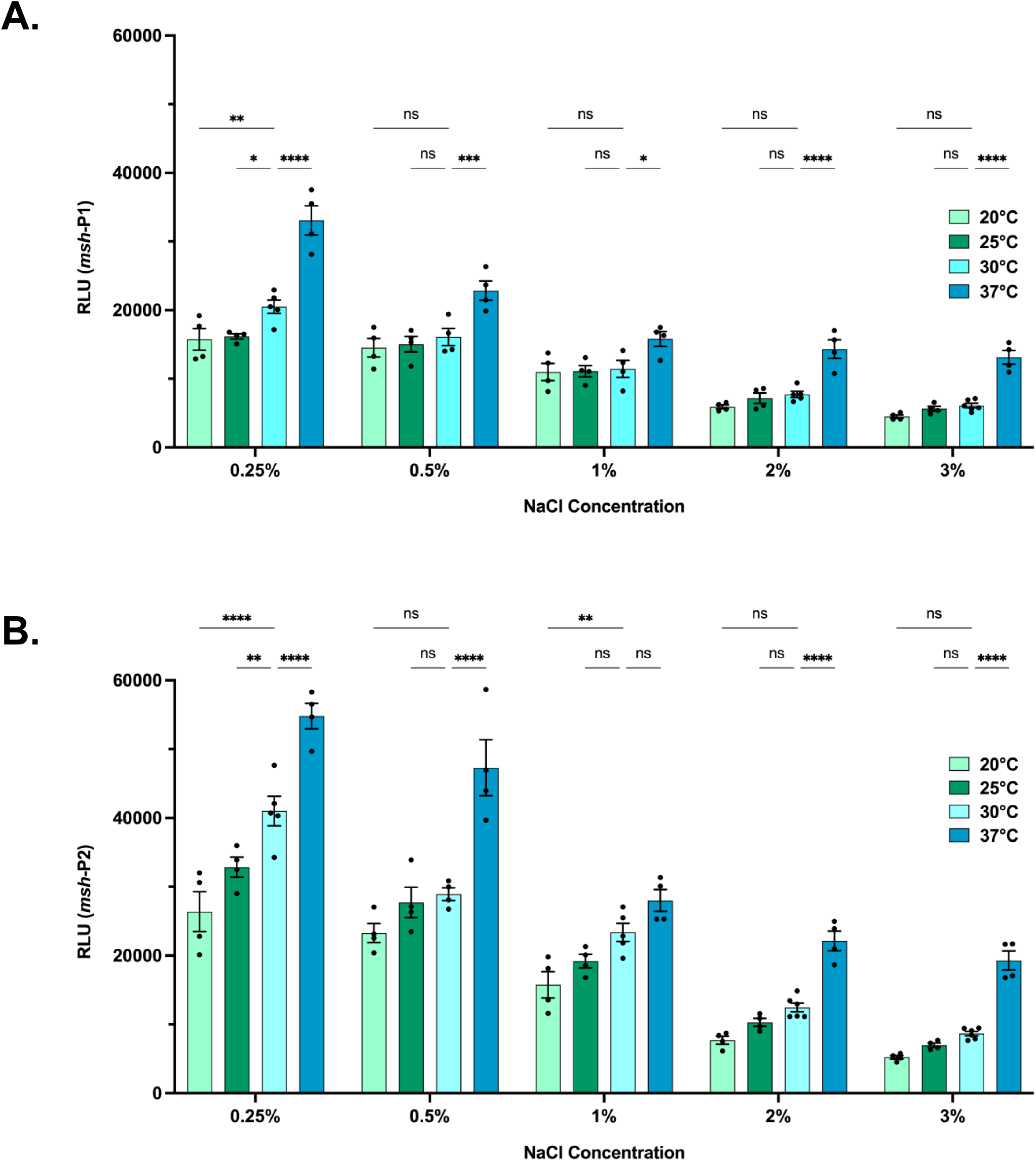
Data presented in Figure 5 plotted by temperature at each salinity. **(A-B)** RLU = relative light units. Data given as mean ± SEM, from a minimum of three biological replicates. **(A)** The promoter activity of *msh*-P1 in mid-exponential phase. Statistical analysis: Two-way ANOVA with Dunnett’s correction, comparing activity at each temperature to 30°C within salinity; [0.25% NaCl: \**P* = 0.0107, \*\**P* = 0.0046, \*\*\*\**P* < 0.0001], [0.5% NaCl: \*\*\**P* = 0.0001], [1% NaCl : \**P =* 0.0150], [2% NaCl: \*\*\*\**P* < 0.0001], [3% NaCl: \*\*\*\**P* < 0.0001], ns = not significant**. (B)** The promoter activity of *msh*-P2 in mid-exponential phase. Statistical analysis: Two-way ANOVA with Dunnett’s correction, comparing activity at each temperature to 30°C within salinity; [0.25% NaCl: \*\**P* = 0.0016, \*\*\*\**P* < 0.0001], [0.5% NaCl: \*\*\*\**P* < 0.0001], [1% NaCl : \*\**P =* 0.0035], [2% NaCl: \*\*\*\**P* < 0.0001], [3% NaCl: \*\*\*\**P* < 0.0001], ns = not significant.

**Figure S3.**
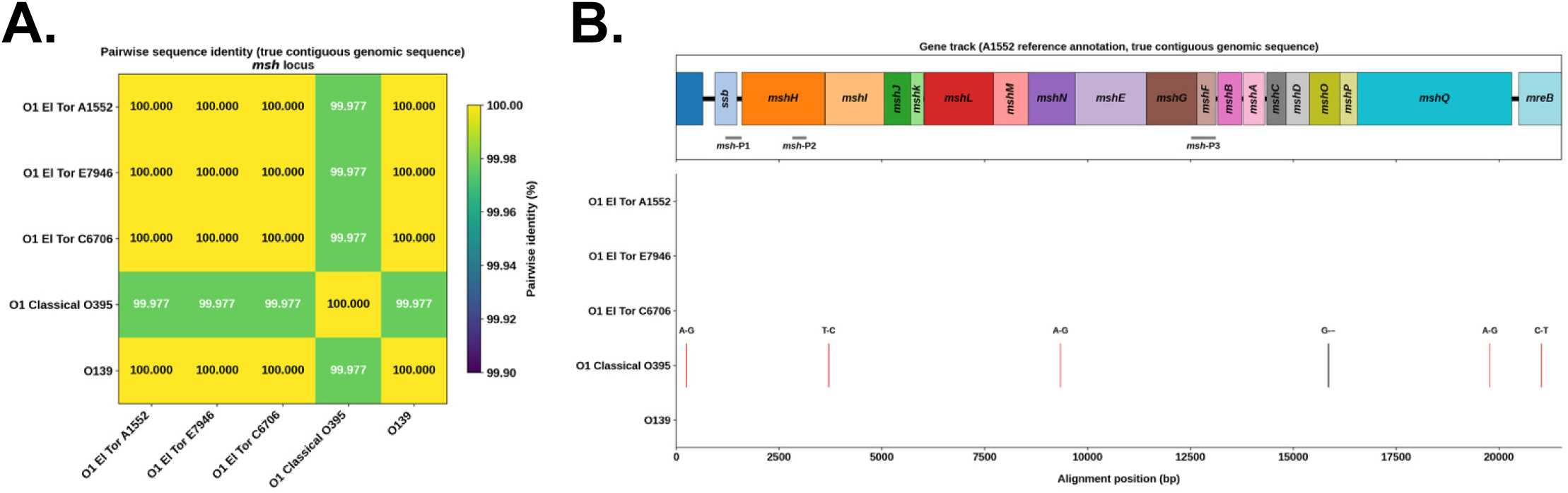
Sequence alignment and variant analysis for the *msh* locus across O1 El Tor strains A1552/E7946/C6706, O1 Classical O395, and O139 serogroups. Contiguous genomic sequences spanning the *msh* locus (*mshH*–*mshQ*, plus flanking *csgD*, *ssb*, and *mreB*) were obtained from complete chromosome assemblies of five *Vibrio cholerae* strains (El Tor A1552, GenBank CP028894.1; El Tor E7946, CP024162.1; El Tor C6706, CP157384.1; Classical O395, CP045719.1; O139, ENA LT992486.1) by extracting the homologous region identified via BLASTn (v2.12.0+, NCBI). Sequences were aligned using MAFFT v7.505 (FFT-NS-2, default settings). Pairwise nucleotide identity was calculated across ungapped alignment columns for each sequence pair. Sliding-window identity (200 bp window, 20 bp step) and per-column variant calls (substitutions and indels relative to the majority-consensus base) were computed in Python (Biopython, NumPy). Figures were generated with Matplotlib. *AI disclosure*: Sequence alignment, identity/variant calculations, and figure generation were performed with the assistance of Claude (Anthropic, Claude Sonnet 5). **(A)** Heatmap comparison of sequence identity across strains. **(B)** A variant map of the differences observed across strains; For O395, vertical red lines indicate a base pair change, vertical black lines indicate a base pair deletion, and the base pair changed is shown. Localization of the *msh* promoter regions used for the transcriptional reporters is indicated by horizontal grey lines.

**Table S1.**
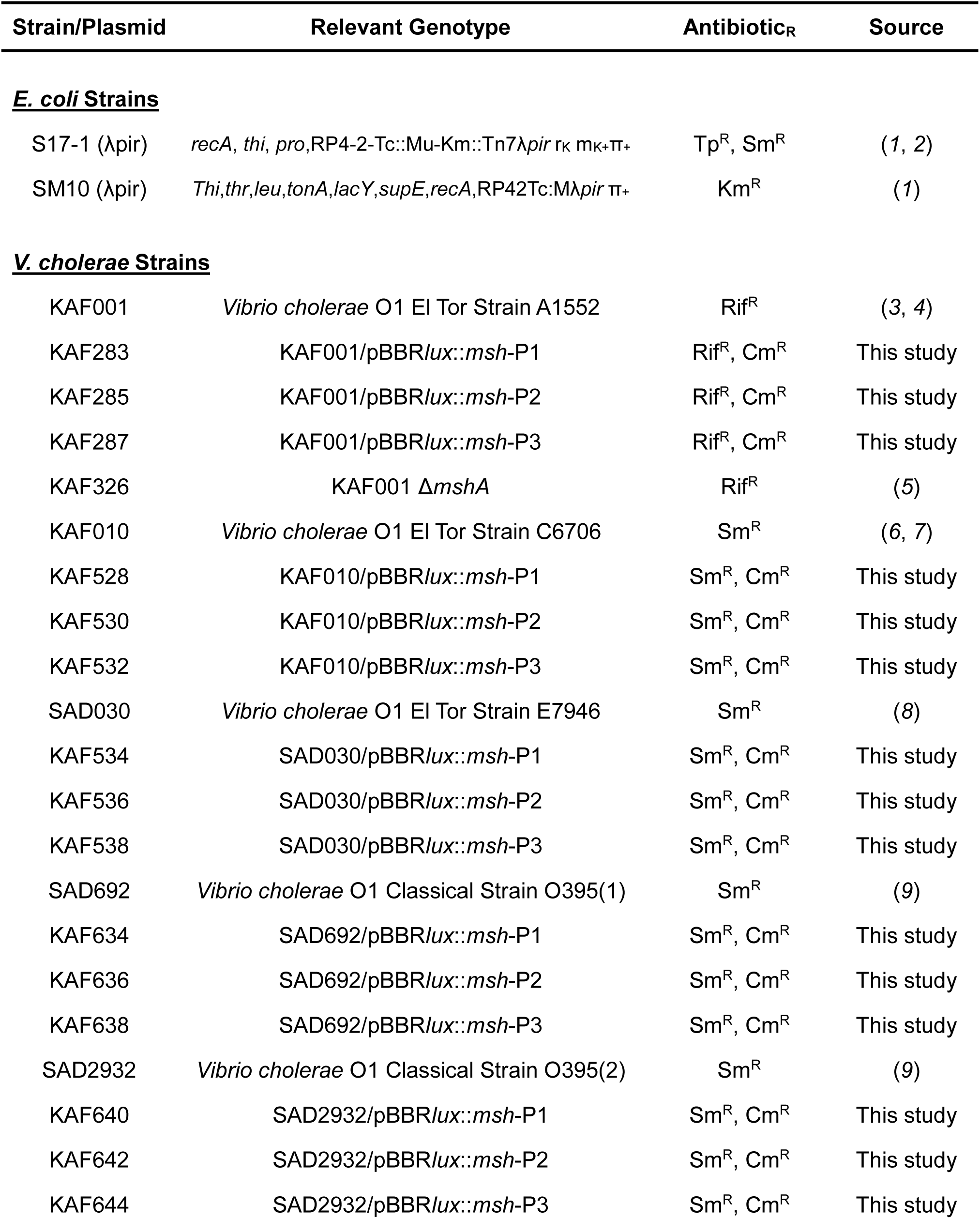
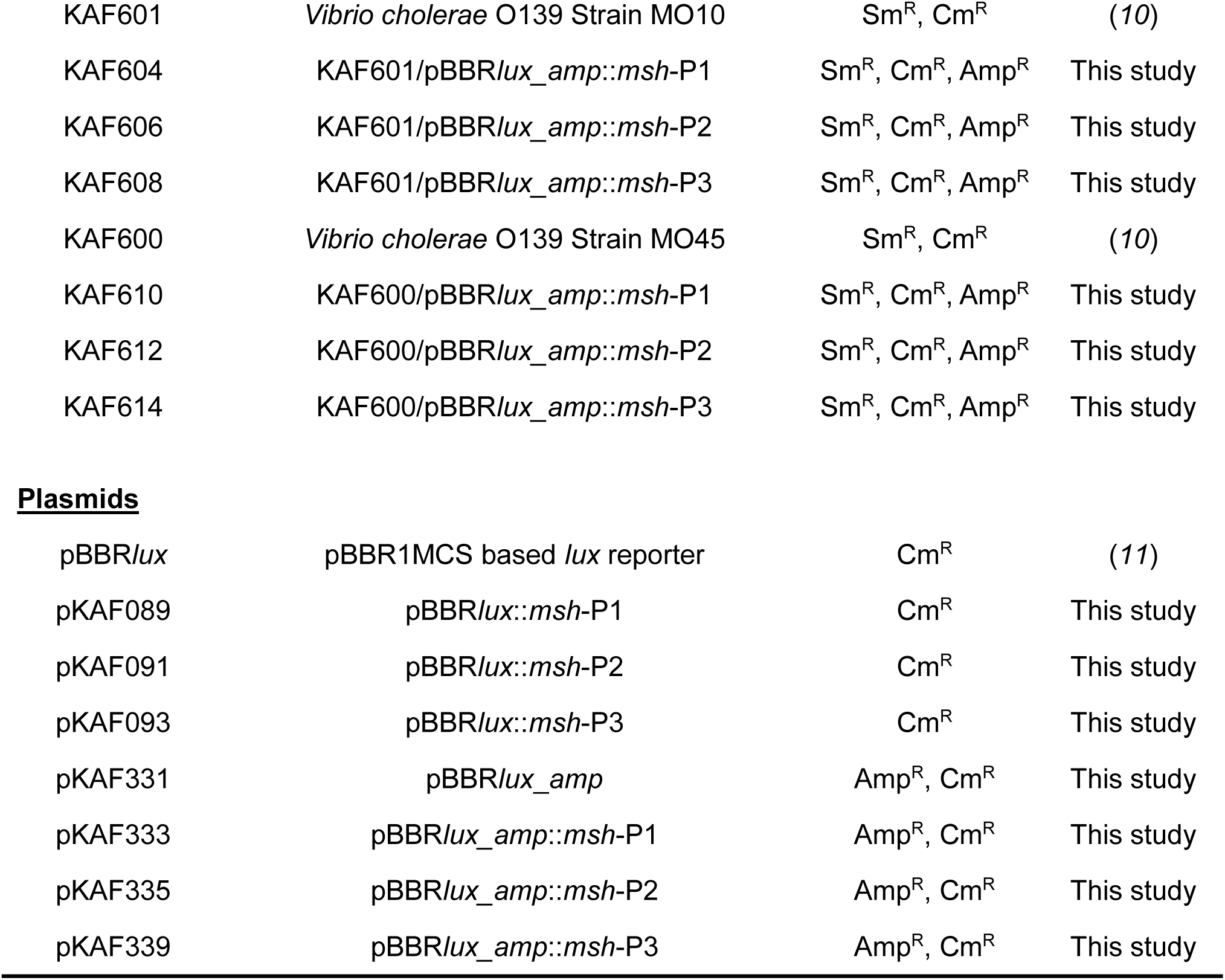
Bacterial Strains and Plasmids used in this study.

| Strain/Plasmid | Relevant Genotype | Antibiotic<sup>R</sup> | Source |
| --- | --- | --- | --- |
| **<u>E. coli Strains</u>** |  |  |  |
| S17-1 (λpir) | *recA, thi, pro*, RP4-2-Tc::Mu-Km::Tn7λpir r<sub>K</sub> m<sub>K</sub>+π<sub>+</sub> | Tp<sup>R</sup>, Sm<sup>R</sup> | (1, 2) |
| SM10 (λpir) | *Thi, thr, leu, tonA, lacY, supE, recA*, RP42Tc:Mλpir π<sub>+</sub> | Km<sup>R</sup> | (1) |
| **<u>V. cholerae Strains</u>** |  |  |  |
| KAF001 | *Vibrio cholerae* O1 El Tor Strain A1552 | Rif<sup>R</sup> | (3, 4) |
| KAF283 | KAF001/pBBRlux::msh-P1 | Rif<sup>R</sup>, Cm<sup>R</sup> | This study |
| KAF285 | KAF001/pBBRlux::msh-P2 | Rif<sup>R</sup>, Cm<sup>R</sup> | This study |
| KAF287 | KAF001/pBBRlux::msh-P3 | Rif<sup>R</sup>, Cm<sup>R</sup> | This study |
| KAF326 | KAF001 Δ*mshA* | Rif<sup>R</sup> | (5) |
| KAF010 | *Vibrio cholerae* O1 El Tor Strain C6706 | Sm<sup>R</sup> | (6, 7) |
| KAF528 | KAF010/pBBRlux::msh-P1 | Sm<sup>R</sup>, Cm<sup>R</sup> | This study |
| KAF530 | KAF010/pBBRlux::msh-P2 | Sm<sup>R</sup>, Cm<sup>R</sup> | This study |
| KAF532 | KAF010/pBBRlux::msh-P3 | Sm<sup>R</sup>, Cm<sup>R</sup> | This study |
| SAD030 | *Vibrio cholerae* O1 El Tor Strain E7946 | Sm<sup>R</sup> | (8) |
| KAF534 | SAD030/pBBRlux::msh-P1 | Sm<sup>R</sup>, Cm<sup>R</sup> | This study |
| KAF536 | SAD030/pBBRlux::msh-P2 | Sm<sup>R</sup>, Cm<sup>R</sup> | This study |
| KAF538 | SAD030/pBBRlux::msh-P3 | Sm<sup>R</sup>, Cm<sup>R</sup> | This study |
| SAD692 | *Vibrio cholerae* O1 Classical Strain O395(1) | Sm<sup>R</sup> | (9) |
| KAF634 | SAD692/pBBRlux::msh-P1 | Sm<sup>R</sup>, Cm<sup>R</sup> | This study |
| KAF636 | SAD692/pBBRlux::msh-P2 | Sm<sup>R</sup>, Cm<sup>R</sup> | This study |
| KAF638 | SAD692/pBBRlux::msh-P3 | Sm<sup>R</sup>, Cm<sup>R</sup> | This study |
| SAD2932 | *Vibrio cholerae* O1 Classical Strain O395(2) | Sm<sup>R</sup> | (9) |
| KAF640 | SAD2932/pBBRlux::msh-P1 | Sm<sup>R</sup>, Cm<sup>R</sup> | This study |
| KAF642 | SAD2932/pBBRlux::msh-P2 | Sm<sup>R</sup>, Cm<sup>R</sup> | This study |
| KAF644 | SAD2932/pBBRlux::msh-P3 | Sm<sup>R</sup>, Cm<sup>R</sup> | This study |
| KAF601 | *Vibrio cholerae* O139 Strain MO10 | Sm<sup>R</sup>, Cm<sup>R</sup> | (10) |
| KAF604 | KAF601/pBBR*lux_amp::msh*-P1 | Sm<sup>R</sup>, Cm<sup>R</sup>, Amp<sup>R</sup> | This study |
| KAF606 | KAF601/pBBR*lux_amp::msh*-P2 | Sm<sup>R</sup>, Cm<sup>R</sup>, Amp<sup>R</sup> | This study |
| KAF608 | KAF601/pBBR*lux_amp::msh*-P3 | Sm<sup>R</sup>, Cm<sup>R</sup>, Amp<sup>R</sup> | This study |
| KAF600 | *Vibrio cholerae* O139 Strain MO45 | Sm<sup>R</sup>, Cm<sup>R</sup> | (10) |
| KAF610 | KAF600/pBBR*lux_amp::msh*-P1 | Sm<sup>R</sup>, Cm<sup>R</sup>, Amp<sup>R</sup> | This study |
| KAF612 | KAF600/pBBR*lux_amp::msh*-P2 | Sm<sup>R</sup>, Cm<sup>R</sup>, Amp<sup>R</sup> | This study |
| KAF614 | KAF600/pBBR*lux_amp::msh*-P3 | Sm<sup>R</sup>, Cm<sup>R</sup>, Amp<sup>R</sup> | This study |
| Plasmids |  |  |  |
| pBBR*lux* | pBBR1MCS based *lux* reporter | Cm<sup>R</sup> | (11) |
| pKAF089 | pBBR*lux::msh*-P1 | Cm<sup>R</sup> | This study |
| pKAF091 | pBBR*lux::msh*-P2 | Cm<sup>R</sup> | This study |
| pKAF093 | pBBR*lux::msh*-P3 | Cm<sup>R</sup> | This study |
| pKAF331 | pBBR*lux_amp* | Amp<sup>R</sup>, Cm<sup>R</sup> | This study |
| pKAF333 | pBBR*lux_amp::msh*-P1 | Amp<sup>R</sup>, Cm<sup>R</sup> | This study |
| pKAF335 | pBBR*lux_amp::msh*-P2 | Amp<sup>R</sup>, Cm<sup>R</sup> | This study |
| pKAF339 | pBBR*lux_amp::msh*-P3 | Amp<sup>R</sup>, Cm<sup>R</sup> | This study |

**Table S2.**
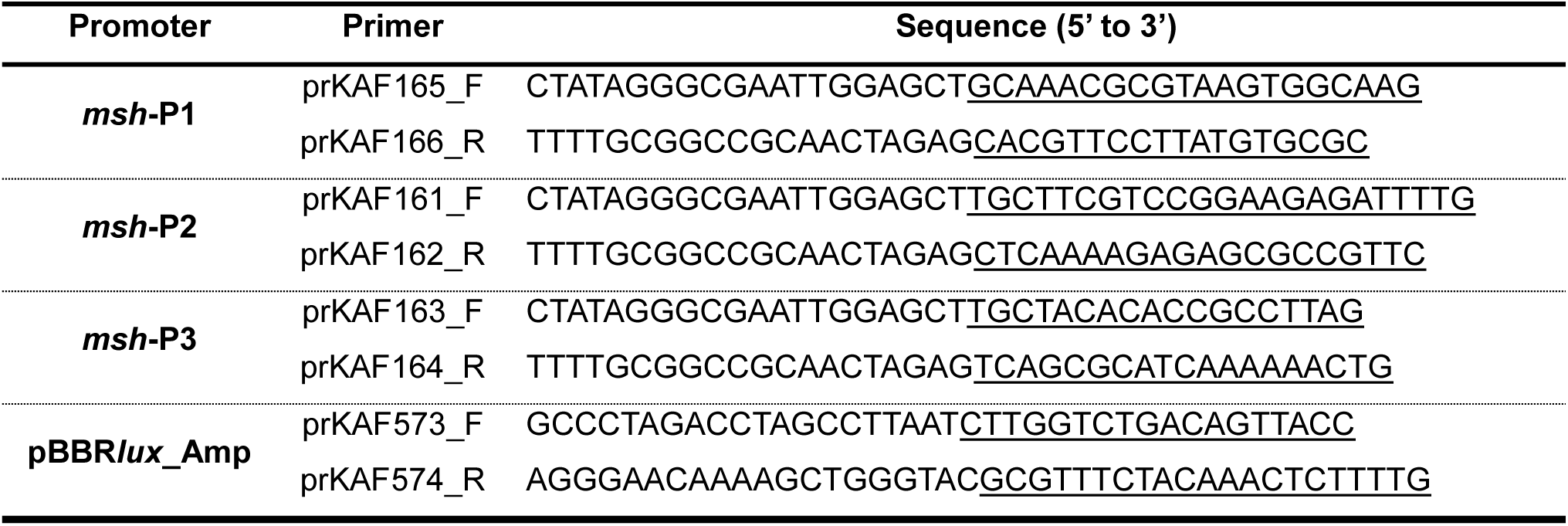
Primer sequences for amplification of promoters used in this study.

